# Spatially Varying Graphical Models for Cell-Cell Interaction Networks in Multiplexed Tissue Imaging

**DOI:** 10.64898/2026.04.01.715977

**Authors:** Sagnik Bhadury, Jeremy T. Gaskins, Arvind Rao

## Abstract

Multiplexed tissue imaging platforms resolve dozens of cell types at single-cell spatial resolution, enabling characterization of conditional interaction networks governing tumor immune microenvironments. Existing methods rely on marginal pairwise co-occurrence statistics without conditioning on third cell types, or estimate a global interaction coefficient per cell type pair that ignores spatial heterogeneity across tissue compartments. We present GP-GHS, a Bayesian nodewise regression framework for inferring spatially varying cell-cell interaction networks from multiplexed imaging data. Each regression coefficient is modeled as a Gaussian process over the tissue domain, approximated via a Hilbert Space Gaussian Process (HSGP) expansion for scalability. A group horseshoe prior assigns a single local shrinkage parameter across all spectral basis coefficients for each candidate edge, enforcing edge inclusion as a group decision rather than independent coefficient level decisions. This separation of roles, where group shrinkage governs edge existence and the spectral prior governs spatial smoothness conditional on existence, enables recovery of spatially structured graphs with high sensitivity. Posterior inference uses a closed form block Gibbs sampler with nodewise regressions parallelized across cores. In simulation studies, GP-GHS dominates all competitors on F1 and MCC across sparsity levels and problem sizes, with ablations isolating group shrinkage as the critical modeling ingredient. Applied to a 140 image CODEX dataset from advanced colorectal cancer patients stratified by pathology subtype, GP-GHS identifies 13 differentially active edges at FDR < 0.05, forming a Treg-centered immunosuppressive network amplified in the diffuse inflammatory subtype, consistent with known mechanisms of Treg recruitment and macrophage-mediated immunosuppression in colorectal cancer.

## 1 Introduction

The tumor microenvironment (TME) is not a passive backdrop to malignant growth but an active, spatially organized ecosystem in which immune, stromal, vascular, and epithelial cell populations engage in continuous, location-dependent crosstalk that shapes immune surveillance, therapeutic resistance, and patient prognosis (Hanahan, 2022; Hinshaw and Shevde, 2019). Recent work for characterizing the TME through bulk and single-cell transcriptomics have established the cellular cast and broad signaling repertoire at play; what has been harder to recover is the spatial structure of these interactions, that is, which cell types conditionally co-localize with which others in specific tissue compartments, and whether those spatial dependencies differ between pathologically distinct disease states. This spatial dimension is not merely descriptive. There is compelling evidence that the geometric organization of immune and stromal cells, the relative positioning of tumor and effector populations, and the spatial fragmentation or coupling of immune neighborhoods are independently predictive of clinical outcome across multiple cancer types (Schürch et al., 2020; Bhadury et al., 2026; Feng et al., 2023).

Multiplexed imaging technologies have made it possible, for the first time, to quantify cell type abundance and spatial location simultaneously across dozens of protein markers in intact tissue sections. Platforms including CODEX (Black et al., 2021), imaging mass cytometry (Giesen et al., 2014), and cyclic immunofluorescence (Lin et al., 2018) now routinely profile 20 to 60 cell types per section at single-cell resolution, generating datasets in which every cell has both a molecular phenotype and a spatial coordinate. These platforms have already yielded biological insights that would have been invisible to dissociation-based approaches: the identification of conserved cellular neighborhoods in colorectal cancer (Schürch et al., 2020), the discovery of spatially distinct immune subtypes in triple-negative breast cancer (Gruosso et al., 2019), and the demonstration that spatial proximity of CD8^+^ T cells to tumor cells is a stronger prognostic marker than CD8^+^ T cell abundance alone (Feng et al., 2023). As these platforms become standard tools in translational oncology, the demand for rigorous statistical methods to extract conditional interaction networks from the resulting spatial data is growing rapidly.

The computational response to this demand has largely followed two tracks that are each, in different ways, insufficient for the problem. The first track consists of cell type cooccurrence and neighborhood enrichment methods, including the widely used approaches implemented in Squidpy (Palla et al., 2022), Giotto (Dries et al., 2021), histoCAT (Schapiro et al., 2017), and related tools. These methods characterize the spatial proximity structure of cell types through permutation tests on neighborhood count matrices or bivariate cooccurrence statistics. They are computationally lightweight and broadly applicable, but they operate on marginal pairwise relationships and have no mechanism for conditional inference. Because they do not control for the presence of third cell types, a co-occurrence signal between, say, macrophages and Tregs can arise spuriously whenever both populations are jointly enriched in the same tissue compartment alongside a third cell type, such as stroma, that recruits both independently. Conditional independence, which is what a graphical model edge actually represents, cannot be recovered from marginal co-occurrence.

The second track consists of methods that explicitly model conditional dependencies among cell types using graphical model machinery. Spatial variance component analysis (SVCA; Arnol et al., 2019) decomposes individual gene and protein expression variability into intrinsic, environmental, and cell-cell interaction components through a Gaussian process random effect model, providing gene-level estimates of how much expression variance is attributable to cellular neighborhood composition. While SVCA was an important methodological advance in framing the spatial interaction problem as a statistical estimation problem rather than a proximity test, it does not recover a network: it assigns a scalar variance fraction to each gene rather than estimating a sparse graph of conditional dependencies among cell types. SpaCeNet (Schrod et al., 2024) extends this idea toward graphical model estimation by modeling distance-dependent pairwise interaction potentials between cell types using conditional independence regularization, but the interaction strengths it infers are global summaries that do not vary across tissue locations. This is a significant limitation, because the TME is not spatially homogeneous: Treg-macrophage interactions that characterize the invasive margin of a tumor may be entirely absent in the tumor core, and a method that pools evidence across all tissue locations to estimate a single global interaction coefficient will systematically mischaracterize the interaction structure in heterogeneous tissues. Informed Spatially Aware Patterns for Multiplexed Immunofluorescence Data (ISPAT; Bhadury et al., 2026) models cell-cell interaction profile varying over tumor trajectory using a gaussian process mixed effect model through conditional independence structure but the covariance estimation relies on a posthoc multi-study factor analytic form.

Separately from the spatial interaction literature, Bayesian graphical models with continuous shrinkage priors have been developed for non-spatial omics data. The graphical horseshoe estimator of Li et al. (2019) places independent half-Cauchy local shrinkage parameters on the off-diagonal elements of the precision matrix, providing adaptive sparse inference for gene co-expression networks. The nodewise regression approach of Meinshausen and Bühlmann (2006) decomposes the *p*-dimensional graphical model problem into *p* parallelizable regression problems, each amenable to standard regularization. These methods have been applied effectively to genetic network inference and multi-omics integration (Ni et al., 2022), but they are designed for stationary multivariate Gaussian data and have no spatial component: the regression coefficient linking two variables is a scalar, not a function of tissue location. Applying them to spatially structured data from multiplexed imaging conflates the spatial average of a heterogeneous interaction field with a global partial correlation, which is neither the quantity of scientific interest nor the quantity needed to recover the true conditional independence graph.

The gap that motivates this work is therefore specific and methodological: there is no existing method that simultaneously (i) estimates a sparse conditional independence graph over cell types from multiplexed imaging data, (ii) allows each edge in that graph to have a spatially varying strength that differs across tissue microenvironments, and (iii) makes edge inclusion a coherent all-or-nothing decision at the graph level rather than an aggregate of independent coefficient-level decisions. Requirements (i) and (ii) together are necessary to recover biologically meaningful conditional interaction networks from spatially heterogeneous TME data. Requirement (iii) is a statistical necessity: when a spatially varying interaction field is represented by a set of basis expansion coefficients, applying independent local shrinkage to each coefficient produces no mechanism for the all-or-nothing edge decision that graphical model selection demands, and the resulting false discovery rate grows with the number of basis functions used to represent the spatial field.

We address this gap through GP-GHS, a Bayesian nodewise regression framework that combines three components. First, we model each spatially varying regression coefficient as a Gaussian process using the Hilbert Space Gaussian Process (HSGP) approximation of Riutort-Mayol et al. (2023), which reduces the 𝒪 (*n*^3^) cost of exact GP inference to 𝒪 (*nm*^2^) matrix operations by representing the spatial field on a spectral basis. Second, we place a group horseshoe prior on the spectral basis coefficients corresponding to each candidate edge, sharing a single local shrinkage parameter across all *m*^2^ basis functions within a block. This group structure enforces the edge-level binary decision: when the group shrinkage parameter *λ*_*j*_ collapses toward zero, the entire interaction field for that cell type pair is simultaneously shrunk to zero across all spatial locations, corresponding to edge absence; when *λ*_*j*_ is large, the spectral prior governing within-group spatial smoothness is free to operate. Third, the *p* nodewise regressions are parallelized and combined by the AND rule to recover a symmetric undirected graph, following the neighborhood selection framework of Meinshausen and Bühlmann (2006).

We apply GP-GHS to the CODEX multiplexed tissue imaging dataset of Schürch et al. (2020), which comprises 140 tissue sections from 35 advanced-stage colorectal cancer (CRC) patients stratified by two pathologically distinct tumor microenvironmental sub-types: Crohn’s-like reaction (CLR) and diffuse inflammatory infiltrate (DII). Colorectal cancer exhibits well-characterized spatial heterogeneity in its immune architecture, with CLR and DII subtypes associated with substantially different prognoses (Schürch et al., 2020) and distinct immune infiltration patterns. The dataset therefore provides an ideal test case for whether GP-GHS can recover spatial interaction networks that differ coherently between defined pathological groups. We evaluate GP-GHS against five competitors through a simulation study spanning two problem sizes and four sparsity levels, using scale-free graphs generated by preferential attachment to reflect the hub-dominated topology characteristic of tumor immune networks (Barabási and Albert, 1999; Chen and Mellman, 2017). We then apply the method to the CRC data to identify differentially active edges between CLR and DII, using a linear mixed effects model on the continuous posterior shrinkage score to account for within-patient correlation across the four images per patient.

The main contributions of this work are as follows. We develop the first Bayesian graphical model framework for spatially varying cell-cell interaction networks in multiplexed tissue imaging, combining a scalable Hilbert space Gaussian process approximation with a group horseshoe prior that enforces graph-level sparsity while borrowing spatial strength across the tissue domain. We demonstrate through simulation that the group structure of the prior is the critical ingredient for recovering spatially structured graphs: replacing the group horseshoe with a standard scalar horseshoe prior causes performance to collapse across all sparsity levels and problem sizes, regardless of all other modeling choices, underscoring that coefficient-level shrinkage is fundamentally insufficient for spatial signal aggregation. We apply the method to a colorectal cancer tissue imaging dataset comprising 140 images from 35 patients and identify a Treg-centered immunosuppressive network that is significantly amplified in the diffuse inflammatory infiltrate subtype relative to the Crohn’s-like response subtype, a finding grounded in the known spatial biology of these two subtypes and recovered with high statistical confidence after patient-level multiple testing correction. Modeling the continuous posterior shrinkage score *κ* through a linear mixed model with patient random effects provides substantially greater power for differential testing, a result with direct practical implications for studies that aim to compare spatial interaction networks across patient populations or experimental conditions.

## 2 Methods

### 2.1 Data Structure and Notation

Let ℒ= { *l*_1_, …, *l*_*n*_ } ⊂ ℝ^2^ denote the set of *n* observed spatial locations (tissue spots) within a single tissue section, with each location *l*_*i*_ = (*x*_*i*_, *y*_*i*_) representing a two-dimensional coordinate in physical tissue space. At each location *l*_*i*_, we observe normalized expression values for *p* cell types, collected in the matrix **Y** ∈ ℝ^*n*×*p*^, where *Y*_*is*_ denotes the expression of cell type *s* at location *l*_*i*_. The goal is to recover a sparse undirected graph 𝒢 = ( 𝒱, ℰ ) over the *p* cell types, where 𝒱 denotes the set of vertices and an edge (*s, s*^′^) ∈ ℰ indicates a spatially structured conditional dependency between cell types *s* and *s*^′^ after accounting for all remaining cell types. Critically, we allow the strength of this dependency to vary continuously across the tissue, a feature that distinguishes the proposed framework from classical graphical models.

### 2.2 Nodewise Spatial Regression Framework

We adopt the neighborhood selection strategy of Meinshausen and Bühlmann (2006), extended to accommodate spatially varying regression coefficients. For each cell type *s* ∈ {1, …, *p*}, we formulate the nodewise regression

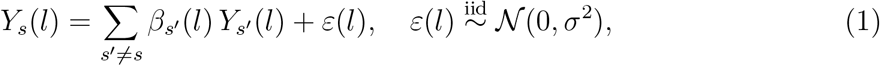

where *Y*_*s*_(*l*) is the expression of cell type *s* at location *l, Y*_*s*_′ (*l*) is the expression of its putative neighbor *s*^′^, and *β*_*s*_′ (*l*) is the spatially varying regression coefficient encoding the conditional association between cell types *s* and *s*^′^ at location *l*. An edge between *s* and *s*^′^ is declared present if the function *β*_*s*_′ ( ·) is not identically zero over the tissue domain. This formulation differs from a standard nodewise lasso in that the coefficients are functions of space rather than scalars, allowing the strength and direction of cell-cell interactions to vary across tissue microenvironments such as the tumor core, invasive margin, and stroma.

The *p* nodewise regressions are independent and are parallelized across cores during computation. A final undirected graph is recovered by applying the AND rule: an edge (*s, s*^′^) is included in ℰ if and only if both the regression of *s* on *s*^′^ and the regression of *s*^′^ on *s* yield an active coefficient function. The AND rule is conservative relative to the OR rule but controls false discoveries more reliably in the moderate-*p* regime typical of multiplexed imaging data (Meinshausen and Bühlmann, 2006).

### 2.3 Spatially Varying Coefficients via Hilbert Space Gaussian Processes

Each spatially varying coefficient *β*_*s*_′ ( · ) is modeled as a realization of a Gaussian process (GP) over the tissue domain. Specifically, we place the prior

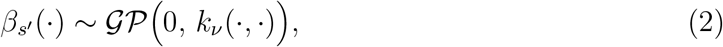

where *k*_*ν*_ is the Matérn covariance kernel with smoothness parameter *ν* and length-scale *ρ*. The Matérn family is preferred over the squared exponential kernel because it allows explicit control over the differentiability of the spatial field: for *ν* = 3*/*2, the resulting process is once mean-square differentiable, which is appropriate for spatial cell-cell interactions that are smooth within tissue compartments but may exhibit sharp transitions at compartment boundaries (Stein, 1999). For two spatial locations *l* and *l*^′^ with Euclidean distance *d* = ∥*l* − *l*^′^∥, the Matérn 3*/*2 kernel takes the form

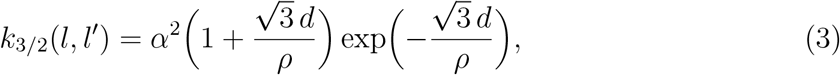

where *α*^2^ is the marginal variance and *ρ* controls the spatial range of dependence. The length-scale *ρ* is set to the median pairwise distance among normalized spatial coordinates by default, though it can be specified by the analyst when prior domain knowledge about interaction range is available.

Exact GP inference requires 𝒪 (*n*^3^) computation due to the Cholesky factorization of the *n* × *n* covariance matrix, which is prohibitive for tissue sections with *n >* 500 spots. We therefore employ the Hilbert Space Gaussian Process (HSGP) approximation of Riutort-Mayol et al. (2023), which approximates the GP using the eigenfunctions of the Laplacian operator on a bounded domain. Specifically, for the two-dimensional domain [ − *L, L*]^2^ obtained by normalizing the spatial coordinates, the Laplacian eigenfunctions factorize as tensor products of one-dimensional sinusoidal basis functions,

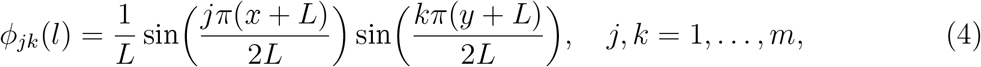

yielding *m*^2^ basis functions in total. The HSGP approximation then represents the spatial field as

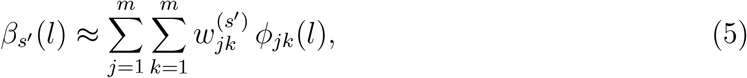

where the basis weights 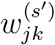 inherit a zero-mean Gaussian prior with variance given by the spectral density of the Matérn kernel evaluated at the corresponding eigenfrequency 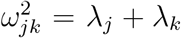 with *λ*_*j*_ = (*jπ/*2*L*)^2^. For the Matérn 3*/*2 kernel in *d* = 2 dimensions, the spectral density is

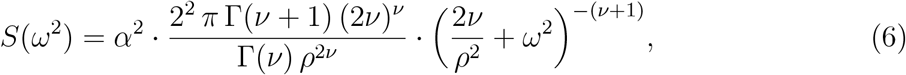

so that 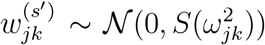. This prior on the basis weights 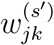 is what encodes spatial smoothness: low-frequency basis functions (small 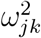) receive large prior variance and are free to capture broad spatial trends, while high-frequency basis functions (large 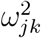 ) are strongly shrunk toward zero, suppressing spatially rough variation. Throughout, we fix *α*^2^ = 1 in *S*(*ω*^2^), as the marginal variance of each interaction field is controlled by the product 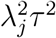 in the group horseshoe hierarchy; absorbing a free *α*^2^ into the prior would create an unidentified redundancy between the GP marginal variance and the group-local shrinkage scale. The HSGP approximation reduces the computational complexity to 𝒪 (*nm*^2^) for a given node regression, making it tractable for the data dimensions encountered in multiplexed imaging.

### 2.4 Group Horseshoe Prior for Edge Selection

The HSGP approximation transforms the spatially varying coefficient model (1) into a linear regression on the expanded design matrix. For node (cell type) *s* with *q* = *p* − 1 neighbors (other cell types), define the blocked design matrix 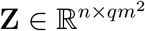 whose *j*-th block of *m*^2^ columns is

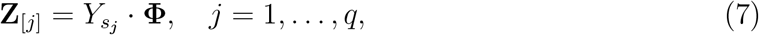

where 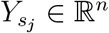 is the expression vector of the *j*-th neighboring cell type and 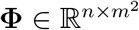 is the matrix of evaluated basis functions. The *j*-th block of regression coefficients 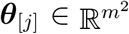 collects the spectral basis weights for neighbor *s*_j_, so that the spatial interaction field is recovered as 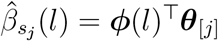.

A critical modeling choice concerns the placement of the sparsity prior within this *qm*^2^-dimensional regression. The naive approach of placing independent local shrinkage parameters on each of the *qm*^2^ predictors - as in a standard scalar horseshoe (Carvalho et al., 2010) - is inappropriate for two reasons that are specific to this problem structure. First, the *m*^2^ basis functions within a block are not scientifically meaningful individually; they are a numerical device for approximating a single spatial function 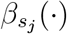 Shrinking individual basis coefficients independently severs this connection and can produce incoherent estimates where a spatial interaction field is partially shrunk across frequency components with no geometric interpretation - for instance, shrinking high-frequency components while retaining low-frequency ones does not correspond to any meaningful biological statement about whether cell type *s*_*j*_ influences *s* in some tissue regions but not others. Second, edge selection in a graphical model is inherently a group-level binary decision: either the entire interaction function 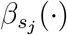 is zero everywhere in the tissue (edge absent) or it is nonzero in at least some region (edge present). An independent scalar prior provides no mechanism for this all-or-nothing decision at the block level and instead accumulates *m*^2^ independent chances to falsely declare activity, inflating false discovery rates in proportion to the number of basis functions keeping in mind that if the variance of *θ*_*jk*_ is not proportional to *S*(*ω*_*k*_) for all *k* within the *j*, then one doesn’t have the gaussian process correspondence.

The appropriate resolution is to share a single local shrinkage parameter *λ*_*j*_ across all *m*^2^ basis coefficients within block *j*, following the grouped regularization framework of Xu et al. (2016). When *λ*_*j*_ → 0, the entire block ***θ***_[_ *j*_]_ is simultaneously collapsed toward zero, so the reconstructed spatial field 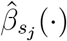 is identically zero across the whole tissue domain - the edge is absent globally. When *λ*_*j*_ is large, all *m*^2^ weights are freed simultaneously, and the within-group spatial pattern is governed entirely by the spectral prior 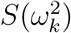. This separation of roles - *λ*_*j*_ controls edge existence, 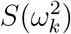 controls spatial smoothness conditional on existence - is the key structural property of the proposed prior and cannot be achieved by any prior that assigns independent local shrinkage to individual basis functions.

Concretely, the *k*-th element of the *j*-th block receives the hierarchical prior

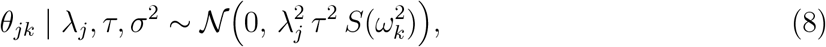

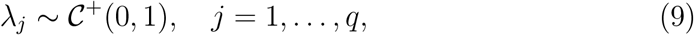

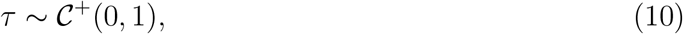

where 𝒞^+^(0, 1) denotes the half-Cauchy distribution (Polson and Scott, 2012). The global parameter *τ* controls the overall graph sparsity - a small *τ* pushes all edges toward absence simultaneously, consistent with the expectation that cell-cell interaction graphs in tissue are sparse. The group-local parameter *λ*_*j*_ controls the existence of individual edge *j* by modulating the variance of the entire *j*-th block through a common scalar. The spectral density 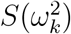 then operates within each surviving group to enforce spatial regularity, giving larger prior variance to low-frequency (smooth) components of 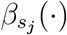 and smaller variance to high-frequency (rough) components. The three levels of the hierarchy thus operate at structurally orthogonal scales: graph-level sparsity, edge-level existence, and within-edge spatial regularity. This parameterization preserves the GP correspondence: the prior variance of *θ*_*jk*_ is 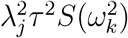, which reduces to the HSGP spectral prior 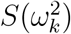 when *λ*_*j*_*τ* = 1. The group structure modulates the overall scale of the entire block through the scalar 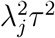, leaving the within-group frequency weighting determined solely by 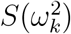. Consequently, when *λ*_*j*_ ^→^ 0 the entire spectral representation collapses proportionally across all frequencies, maintaining the GP structure of the surviving field rather than distorting it.

The group structure also has a direct consequence for how information is accumulated across spatial locations. When evaluating whether edge *j* is active, the posterior for *λ*_*j*_ integrates evidence from all *n* observations and all *m*^2^ basis coefficients jointly through the GP-weighted quadratic form 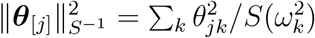, which appears in the full conditional for 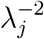 (Section 2.5). A spatially smooth but globally weak interaction - one where 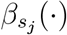 is nonzero but small everywhere - will accumulate sufficient signal across all basis components to prevent *λ*_*j*_ from collapsing, whereas a signal that is present only in isolated individual basis coefficients will be shrunk under the group prior since 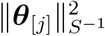 aggregates over the full block. This aggregation property is precisely what distinguishes a group prior from *m*^2^ independent scalar priors and is the mechanism that gives the proposed method its sensitivity advantage over methods that ignore spatial structure entirely.

The group shrinkage factor for edge *j* is defined as 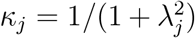, which lies in (0, 1). When *κ*_*j*_ ≈ 1 the group is fully shrunk toward zero and the edge is absent; when *κ*_*j*_ ≈ 0 the group is unshrunk and the edge is present. We declare edge (*s, s*_*j*_) active if the posterior mean 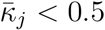, the threshold at which the posterior assigns equal probability to signal and noise for that group (Carvalho et al., 2010).

The likelihood for node *s* is

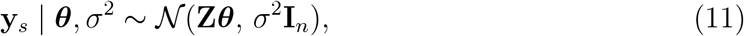

and *σ*^2^ receives the sparse-IG prior *p*(*σ*^2^) ∼ *IG*(0.1, 0.1), which provides mild regularization while remaining diffuse over a wide range of residual variances.

### 2.5 Posterior Inference via Gibbs Sampling

All full conditionals are available in closed form, yielding an efficient block Gibbs sampler. Introducing the auxiliary variables *ξ* and *ν*_*j*_ for the half-Cauchy representations 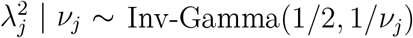 and *τ*^2^| *ξ* ∼ Inv-Gamma(1 / 2 / 1 / *ξ*), the sampler cycles through the following updates.

#### Update θ

The full conditional is multivariate Gaussian,

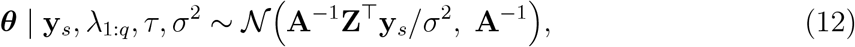

where **A** = **Z**^⊤^**Z***/σ*^2^+**D**^−1^ where **D** is block diagonal with *j*^*th*^ block being 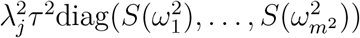, the block-diagonal prior covariance, with the spectral densities determining the within-group scaling.

#### Update λ_j_

For each *j* = 1, …, *q*, the full conditional is

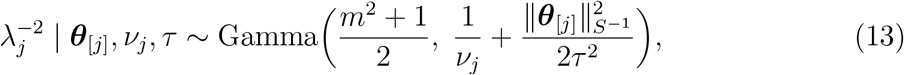

where 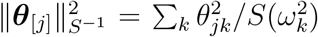 is the GP-adjusted quadratic form. This weighted norm accounts for the spectral structure: coefficients associated with low-frequency (smooth) basis functions are penalized less than those at high frequencies.

#### Update τ

The global shrinkage parameter satisfies

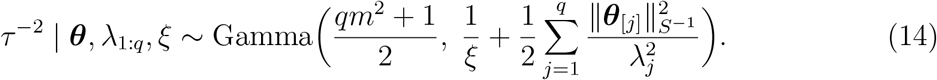

#### Update σ^2^

The residual variance update is

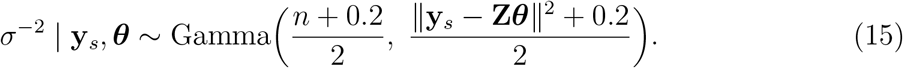

Auxiliary variables *ξ* and *ν*_*j*_ are updated as *ξ*^−1^ | *τ* ^2^ ∼ Gamma(1, 1 + 1*/τ* ^2^) and 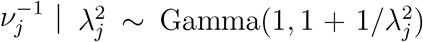. We run the sampler for 3,000 iterations with a burn-in of 1,000 and thinning of 5, retaining 400 posterior draws per node regression. The *p* nodewise samplers are embarrassingly parallel and are distributed across available CPU cores.

### 2.6 Spatial Edge Maps and Posterior Summaries

For each declared active edge (*s, s*^′^), the posterior mean spatial interaction field is recovered as

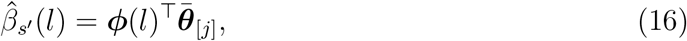

where 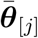 is the posterior mean of the *j*-th coefficient block. This produces a continuous map over the tissue domain assigning a signed interaction strength at every observed location. Regions where 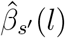 is large and positive indicate spatial co-enrichment of cell types *s* and *s*^′^, while negative values indicate spatial exclusion. Pointwise 95% posterior credible intervals are computed from the posterior samples, providing uncertainty quantification for the spatial maps. The full collection of *p*(*p* − 1)*/*2 edge maps, together with the global adjacency matrix and group-level shrinkage summaries, constitute the output of the proposed method.

#### Graph construction

The *p* nodewise regressions are fit independently, each producing a directed edge list: node *s* declares neighbor *s*^′^ active if 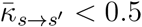, where 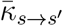, denotes the posterior mean group shrinkage factor from the regression of *s* on { *s*^′^ : *s*^′^ ≠ *s* }. Because the nodewise regressions are not jointly constrained, the directed edge sets need not be symmetric: it is possible that the regression of *s* on *s*^′^ declares the edge active while the regression of *s*^′^ on *s* does not. We resolve this asymmetry by the AND rule: an undirected edge (*s, s*^′^) is included in the estimated graph 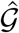 if and only if both regressions declare it active,

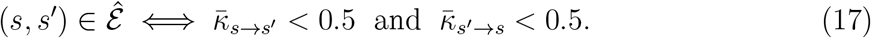

The AND rule is conservative relative to the OR rule (which would include an edge if either regression declares it active) and is known to provide better false discovery control in the moderate-*p* regime (Meinshausen and Bühlmann, 2006). The resulting adjacency matrix ***Â*** ∈ {0, 1}^*p*×*p*^ is symmetric by construction with *Â*_*ss*_′ = *Â*_*s*_′_*s*_ for all pairs. The edge strength for a retained edge is summarized by 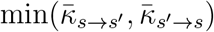, taking the more confident of the two directional posteriors as the representative shrinkage score. *κ* near zero indicates an active signal and *κ* near one indicates shrinkage toward the null. Taking the minimum selects the directional posterior that expresses the stronger evidence of interaction, which is the appropriate summary for a symmetric undirected edge. The full collection of *p*(*p* −1)*/*2 spatial interaction maps for active edges, the symmetric adjacency matrix ***Â***, and the matrix of posterior shrinkage scores constitute the complete output of the method. We present a full schematic overview of the method in Figure 1.

**Figure 1.**
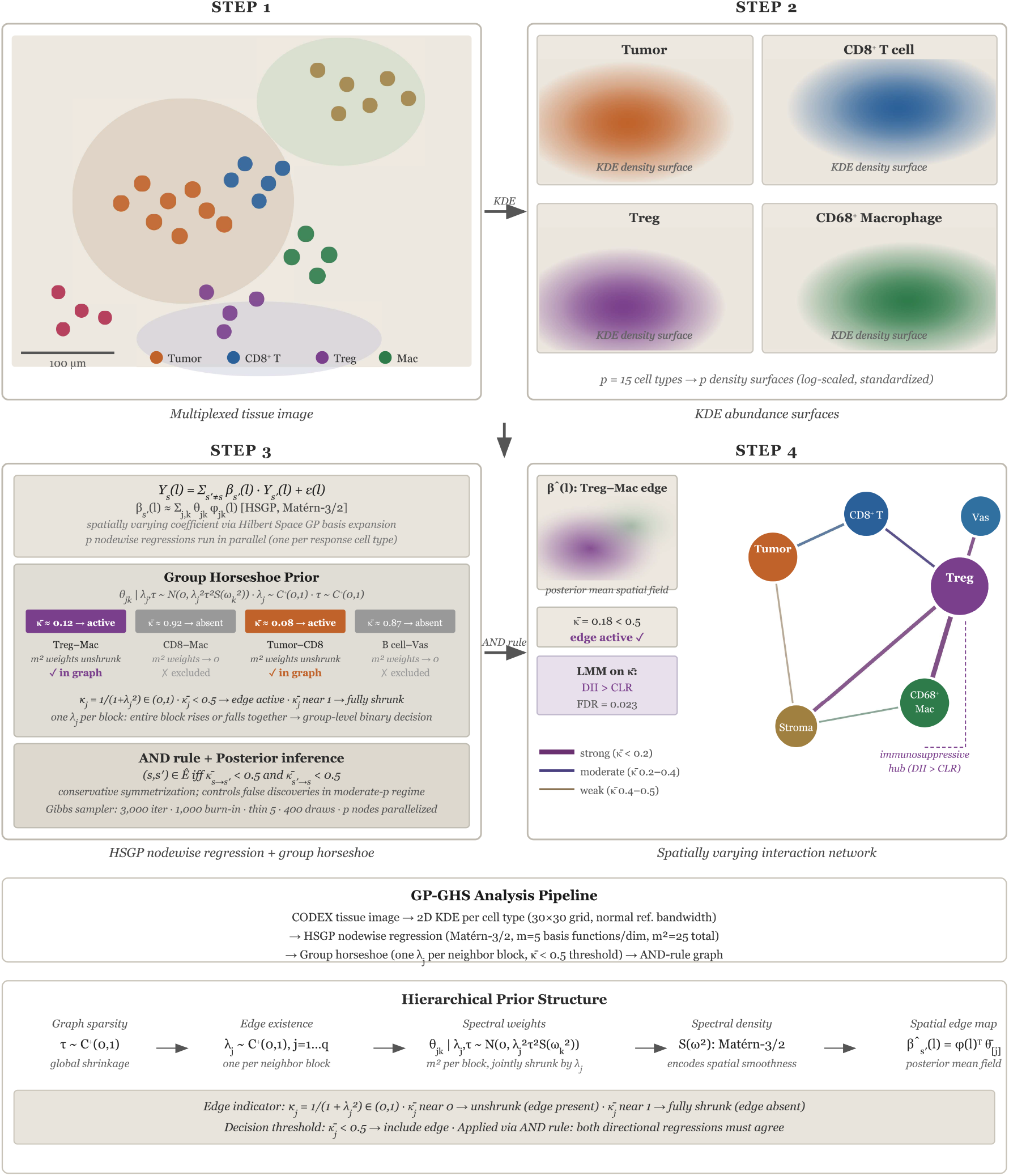
Overview of the GP-GHS pipeline. (**Step 1**) Multiplexed CODEX tissue image with *p* = 15 phenotyped cell types at single-cell spatial resolution. (**Step 2**) Per-cell-type kernel density surfaces on a 30 × 30 grid serve as spatially referenced expression inputs. (**Step 3**) HSGP nodewise regression with a group horseshoe prior: one shrinkage parameter *λ*_*j*_ per neighbor block enforces edge inclusion as a group-level decision via 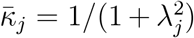, with 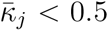 declaring an edge active. (**Step 4**) Inferred spatially varying network with posterior edge maps 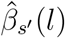 differential testing via linear mixed model on 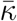 recovers six FDR *<* 0.05 edges between CLR and DII pathology groups.

## 3 Simulation Study

### 3.1 Data Generating Mechanism

We evaluate the proposed method against four competitors through a structured simulation study designed to reflect the key features of multiplexed tissue imaging data. We consider a fixed set of *n* = 600 spatial locations drawn uniformly from the unit square [0, 10]^2^, with *p* = 15, 25 cell types. The true cell-cell interaction graph is generated using the Barabási-Albert preferential attachment model (Barabási and Albert, 1999), which produces a scale-free topology with a small number of hub cell types having many connections and many cell types having few connections. This architecture is biologically motivated: in tumor microenvironments, a handful of central cell types such as macrophages and tumor cells tend to coordinate interactions with many others, while more peripheral populations such as plasma cells or mast cells participate in fewer interactions (Chen and Mellman, 2017). We examine four sparsity levels by targeting edge probabilities of 0.7, 0.5, 0.3, and 0.1, corresponding to dense, moderate, sparse, and very sparse true graphs.

Conditional on the true adjacency matrix **A**^∗^ ∈ { 0, 1 } ^*p*×*p*^, data are generated according to the structural equation

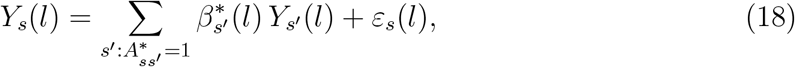

where 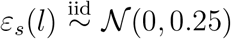 and the true spatially varying coefficients 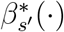 are drawn from a zero-mean GP with Matérn 3 */*2 kernel, appropriate length-scale and marginal standard deviation *α*^∗^ = 0.5. This choice of *ρ*^∗^ relative to the normalized domain produces spatial patterns that vary over a moderate fraction of the tissue extent, consistent with the scale of immune infiltration patterns observed in multiplexed imaging data. Each simulation replicate draws fresh GP realizations for all true edges and fresh noise, ensuring that results reflect variability across both spatial configurations and noise realizations. Ten independent replicates are generated per sparsity level, yielding 40 simulated datasets in total.

### 3.2 Competitor Methods

We compare GP-GHS against five competitor methods spanning penalized likelihood, Bayesian scalar shrinkage, exact GP regression, and naive correlation thresholding.

#### Graphical Lasso (GLasso)

The graphical lasso (Friedman et al., 2008) estimates a sparse precision matrix by solving 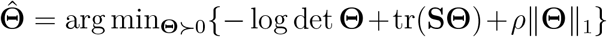, where **S** is the *p* × *p* sample covariance matrix of (*Y*_1_(*l*), *Y*_2_(*l*), …, *Y*_*p*_(*l*)) and *ρ >* 0 is a penalty parameter selected via BIC over a grid of 15 values. GLasso assumes stationarity and ignores spatial coordinates entirely, producing a single global precision matrix.

#### Nodewise Lasso (NL)

Following Meinshausen and Bühlmann (2006), we fit an *ℓ*_1_-penalized regression for each node against all remaining nodes, with the penalty selected by fivefold cross-validation. The undirected graph is recovered via the AND rule. Like GLasso, NL treats all regression coefficients as scalars and is blind to spatial structure.

#### Standard Horseshoe (SHS)

This method applies the scalar horseshoe prior of Carvalho et al. (2010) to each nodewise regression, assigning an independent local shrinkage parameter *λ*_*jk*_ to every scalar predictor rather than one *λ*_*j*_ per neighbor block. Because shrinkage operates at the coefficient level rather than the group level, SHS has no mechanism for coherent all-or-nothing edge decisions: individual basis coefficients within a neighbor block can be selectively shrunk, yielding interaction field estimates that are neither uniformly zero nor uniformly nonzero. Edge inclusion is assessed via the pseudo-*p*-value criterion 2 min { *P* (*θ >* 0 | **y**), *P* (*θ <* 0 | **y**) } *<* 0.05 applied to each scalar coefficient. SHS retains the nodewise Bayesian framework of GP-GHS but discards both the group structure and the GP spectral prior, thereby isolating their joint contribution to performance.

#### Pairwise GP Regression (Exact GP –EGP)

This competitor directly fits an independent Gaussian process regression for each ordered node pair (*s, s*^′^), using a Matérn-3*/*2 kernel with length-scale matched to the HSGP auto-selected *ρ*. For each response node *s*, the partial residual with respect to neighbor *s*^′^ is formed by regressing out all other neighbors via OLS, and the resulting residual is treated as a noisy observation of *β*_*st*_(*l*)*Y*_*t*_(*l*). We term this competitor pairwise because it cannot simultaneously fit GP priors for all *p* − 1 neighbors of a given node. Doing so would require inverting a joint GP covariance matrix of dimension (*p* − 1)*n* × (*p* − 1)*n*, which is computationally prohibitive at the problem sizes considered. The pairwise approach instead uses one-step OLS residuals to partial out the remaining neighbors, replacing a joint GP fit with a sequence of marginal GP fits at substantially lower cost. Edge inclusion is determined by majority vote: an edge (*s, s*^′^) is declared active if the pointwise 95% posterior credible interval excludes zero at more than 50% of spatial locations, and the final undirected graph is symmetrized by the AND rule. EGP is the natural exact baseline for GP-GHS: it shares the same spatial prior and edge selection logic but forgoes the HSGP approximation, group horseshoe shrinkage, and joint multi-neighbor estimation, at substantially greater computational cost.

#### Correlation Threshold (CT)

A naive baseline that declares an edge whenever | *r*_*ss*_′ | *>* 0.15 in the sample correlation matrix. CT makes no structural assumptions and serves as a lower performance bound.

GLasso and NL represent standard penalized likelihood methods for graphical model selection. SHS isolates the added value of the spatial GP component by holding the horseshoe framework fixed while removing spatial structure. EGP isolates the value of the HSGP approximation and group shrinkage by retaining spatial structure while removing them. CT provides a naive lower bound. All methods are applied to the same scaled expression matrices. MCMC-based methods (GP-GHS and SHS) use 2,000 iterations with 500 burn-in and thinning of 5.

### 3.3 Evaluation Metrics

Let ***Â*** ∈ {0, 1}^*p*×*p*^ denote the estimated adjacency matrix and **A**^∗^ the true adjacency matrix. Evaluation is conducted on the upper triangle of both matrices, treating edge recovery as a binary classification problem over the *p*(*p* − 1)*/*2 possible edges for *p* = 15, 25. We compute the following metrics.

Let TP, FP, TN, FN denote true positives, false positives, true negatives, and false negatives respectively. The metrics are:

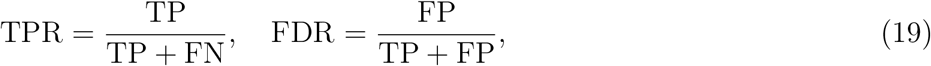

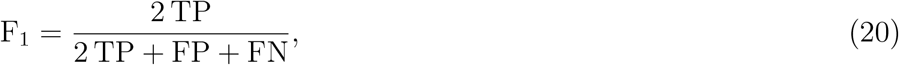

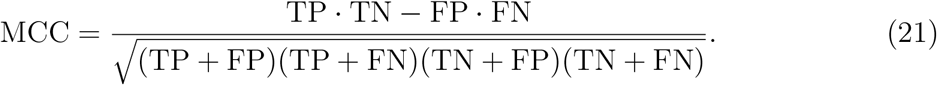

The Matthews Correlation Coefficient (MCC) is particularly informative under class imbalance (Chicco and Jurman, 2020), which arises in the very sparse setting where true edges constitute only ∼ 10% of all pairs. In the same spirit, the F1 score favors methods to select more true positive without making either type I or type II errors. All metrics are averaged across the 10 replicates per sparsity level, and we report means with standard deviations. Wall-clock runtime per dataset is also recorded.

### 3.4 Results: *n* = 600, *p* = 15

Results for the *n* = 600, *p* = 15 configuration are presented in Figure 2. We organize the discussion around five substantive findings that together characterize the operating properties of GP-GHS and the newly added Exact GP competitor relative to the remaining methods.

**Figure 2.**
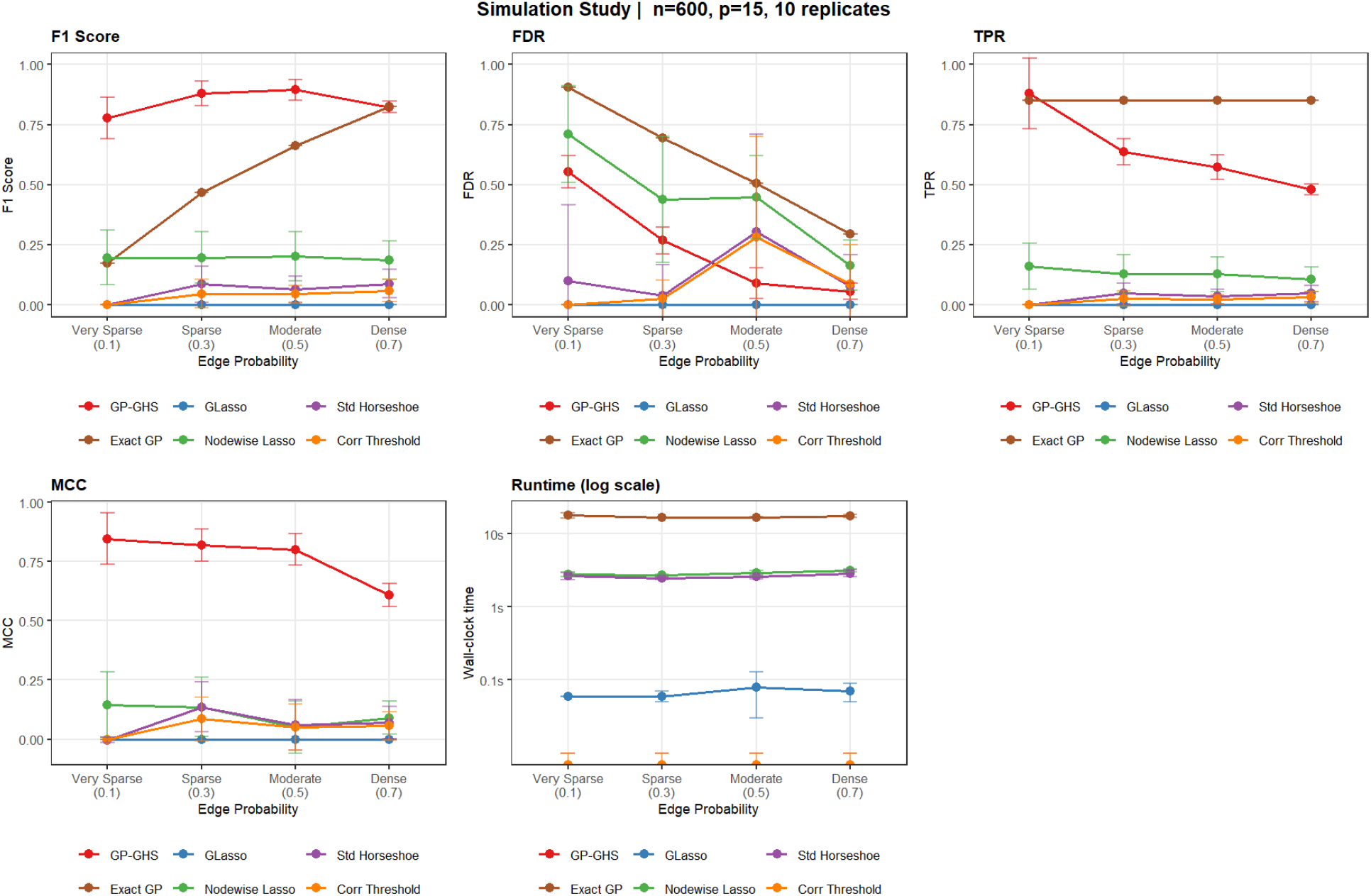
Simulation results for *n* = 600, *p* = 15, scale-free true graph, 10 replicates per sparsity level. Each panel shows mean ± one standard deviation across replicates. Edge probability on the horizontal axis corresponds to expected edge counts of approximately 11 (very sparse, *π* = 0.1), 32 (sparse, *π* = 0.3), 53 (moderate, *π* = 0.5), and 74 (dense, *π* = 0.7) out of 105 possible edges. The runtime panel uses a log_10_ scale.

#### Finding 1: GP-GHS dominates all competitors on F_1_ and MCC across the full sparsity range, with Exact GP competitive only in dense graphs

GP-GHS achieves F_1_ scores of approximately 0.78, 0.88, 0.88, and 0.83 at edge probabilities *π* ∈ { 0.1, 0.3, 0.5, 0.7 } respectively, with corresponding MCC values of 0.83, 0.80, 0.78, and 0.62. The Exact GP competitor displays a qualitatively different trajectory: its F_1_ rises monotonically from near zero at *π* = 0.1 to approximately 0.83 at *π* = 0.7, effectively matching GP-GHS only in the densest configuration. In all sparser settings Exact GP substantially underperforms GP-GHS on F_1_ and MCC despite sharing the same Matérn spatial prior. All remaining methods (GLasso, Nodewise Lasso, Standard Horseshoe, Correlation Threshold) stay below F_1_ ≈ 0.20 at every sparsity level, confirming that neither penalized likelihood nor scalar Bayesian shrinkage is adequate for spatially structured graph recovery.

The performance of GP-GHS peaks in the sparse-to-moderate range (*π* ∈ { 0.3, 0.5 } ) and declines modestly at the dense extreme, a pattern consistent with the behavior of the global horseshoe parameter *τ* . When the graph is moderately sparse, *τ* finds a well-separated operating point that discriminates active from inactive neighbor blocks. At *π* = 0.7, with roughly 74 true edges out of 105, the prior diffuses its shrinkage mass across a large number of active blocks, reducing fine-grained discrimination and producing the observed decline in MCC from 0.83 to 0.62.

#### Finding 2: Exact GP achieves high sensitivity throughout but suffers severe false discovery inflation in sparse graphs

The TPR of Exact GP is remarkably stable across sparsity levels, remaining near 0.85 from *π* = 0.1 through *π* = 0.7. This is the highest TPR of any method in moderate and dense settings and matches GP-GHS at very sparse and sparse settings. However, its FDR is catastrophically high in sparse configurations: approximately 0.90 at *π* = 0.1 and 0.70 at *π* = 0.3, meaning the vast majority of declared edges are false positives. The FDR of Exact GP improves as the graph densifies, reaching approximately 0.30 at *π* = 0.7, but remains substantially above GP-GHS at every setting.

This behavior reflects a structural property of the Exact GP edge selection rule. The majority-vote criterion — declaring an edge active if the pointwise 95% credible interval excludes zero at more than 50% of locations — treats each spatial pair independently and has no global regularization across edges. In a sparse graph, the marginal posterior for each node pair integrates only the local residual signal for that pair, without borrowing the information that most pairs should be null. Consequently, the posterior credible intervals for noise-only pairs frequently exclude zero at a nontrivial fraction of locations due to spatially coherent noise, producing widespread false inclusions. GP-GHS avoids this by placing a group horseshoe prior on the entire collection of neighbor blocks jointly, so that the global shrinkage parameter *τ* calibrates edge-level sparsity in a data-adaptive manner that reflects the overall graph density. The FDR contrast between the two spatial methods at *π* = 0.1 — approximately 0.90 for Exact GP versus 0.55 for GP-GHS — directly quantifies the value of this joint sparsity regularization.

#### Finding 3: GP-GHS maintains high sensitivity and controls FDR progressively across sparsity levels

GP-GHS achieves TPR of approximately 0.85, 0.85, 0.57, and 0.48 across the four sparsity levels, consistently the highest among methods that also maintain reasonable FDR. The FDR of GP-GHS declines monotonically from ≈ 0.55 at *π* = 0.1 to ≈ 0.05 at *π* = 0.7, indicating that precision improves as the graph densifies and the horseshoe hierarchy concentrates its mass more uniformly on active edges. The elevated FDR at *π* = 0.1 is expected: at this sparsity level only roughly 11 true edges exist, and the global shrinkage parameter must simultaneously suppress approximately 94 null pairs while protecting a small number of true signals, a regime that inevitably trades some precision for recall. The large standard deviations on GP-GHS FDR at *π* = 0.1 (spanning roughly 0.30 to 0.80 across replicates) reflect genuine replicate-to-replicate variability driven by the small number of true signals rather than instability of the estimator.

#### Finding 4: SHS and GLasso confirm that spatial structure and group shrink-age are jointly necessary

Standard Horseshoe (SHS) shares the nodewise Bayesian framework with GP-GHS but discards the group prior and GP spectral prior in favor of independent scalar shrinkage on each basis coefficient. Its F_1_ remains below 0.05 and its TPR near zero across all sparsity levels, performance indistinguishable from GLasso and Correlation Threshold. The near-complete failure of SHS is informative: it establishes that the Bayesian nodewise structure alone confers no advantage over standard penalized methods when spatial structure is ignored. The critical contributions of GP-GHS are the group horseshoe hierarchy, which aggregates evidence across all *m*^2^ spectral basis coefficients to make each edge inclusion a joint decision, and the GP spectral prior, which encodes the expectation that true interaction fields are spatially smooth and therefore generate coherent signal accumulation across basis frequencies. Without both components, the horseshoe prior has no mechanism to distinguish a weak but spatially structured edge from *m*^2^ independent noise fluctuations.

GLasso performs uniformly poorly (F_1_ *<* 0.10 throughout) for a different reason: it estimates a single stationary precision matrix from 600 spatially varying observations, conflating the spatial average of each interaction field with its global marginal association. The *ℓ*_1_ penalty then operates on this averaged signal rather than the spatially resolved interaction, selecting edges based on marginal correlation structure that is unrelated to the true conditional dependence network. This is not a limitation of the graphical lasso as a general tool but rather a demonstration that methods designed for exchangeable observations cannot be applied to spatially heterogeneous data without fundamental modification.

#### Finding 5: GP-GHS achieves its performance at a computational cost that remains tractable, while Exact GP is the most expensive competitor

GP-GHS has a mean wall-clock time of approximately 2–3 seconds per dataset in these experiments, which reflects 11-core parallelization of *p* = 15 nodewise regressions with *m* = 4 basis functions per dimension (*m*^2^ = 16 spectral components) and NMC = 2,000 iterations. The HSGP approximation reduces the per-iteration cost from 𝒪 (*n*^3^) – which would be required for exact GP inference — to 𝒪 (*nm*^2^) matrix products, making the sampler feasible at *n* = 600. Runtime is flat across sparsity levels as expected, since the computational bottleneck is the dimension of the spectral basis system rather than the true graph density.

Exact GP is the most expensive method at approximately 10–20 seconds per dataset, roughly 5–8 × slower than GP-GHS despite being run sequentially rather than in parallel. Its cost grows as 𝒪 (*p*(*p* − 1)*n*^3^) due to the Cholesky factorization required per node-neighbor pair, and the near-constant runtime across sparsity levels reflects the fact that all *p*(*p* − 1) node pairs are evaluated regardless of the true graph. Nodewise Lasso and SHS occupy an intermediate range of approximately 2–3 seconds, while GLasso is the fastest method at roughly 0.1 seconds due to its closed-form coordinate descent updates. Correlation Threshold is effectively instantaneous. The computational advantage of GP-GHS over Exact GP – combined with its substantially better FDR control – establishes that the HSGP approximation and group horseshoe prior together dominate the exact GP baseline both statistically and computationally.

### 3.5 Results: *n* = 600, *p* = 25

Results for the higher-dimensional configuration (*n* = 600, *p* = 25, 300 possible edges) are presented in Figure 3. The *p* = 25 setting expands the nodewise regression dimension from *q* = 14 to *q* = 24 predictors per response and increases the candidate edge set from 105 to 300 pairs. The central question is whether the performance advantage of GP-GHS documented in Section 3.4 survives the transition to a harder problem, and whether the relative behavior of Exact GP changes qualitatively with dimension.

**Figure 3.**
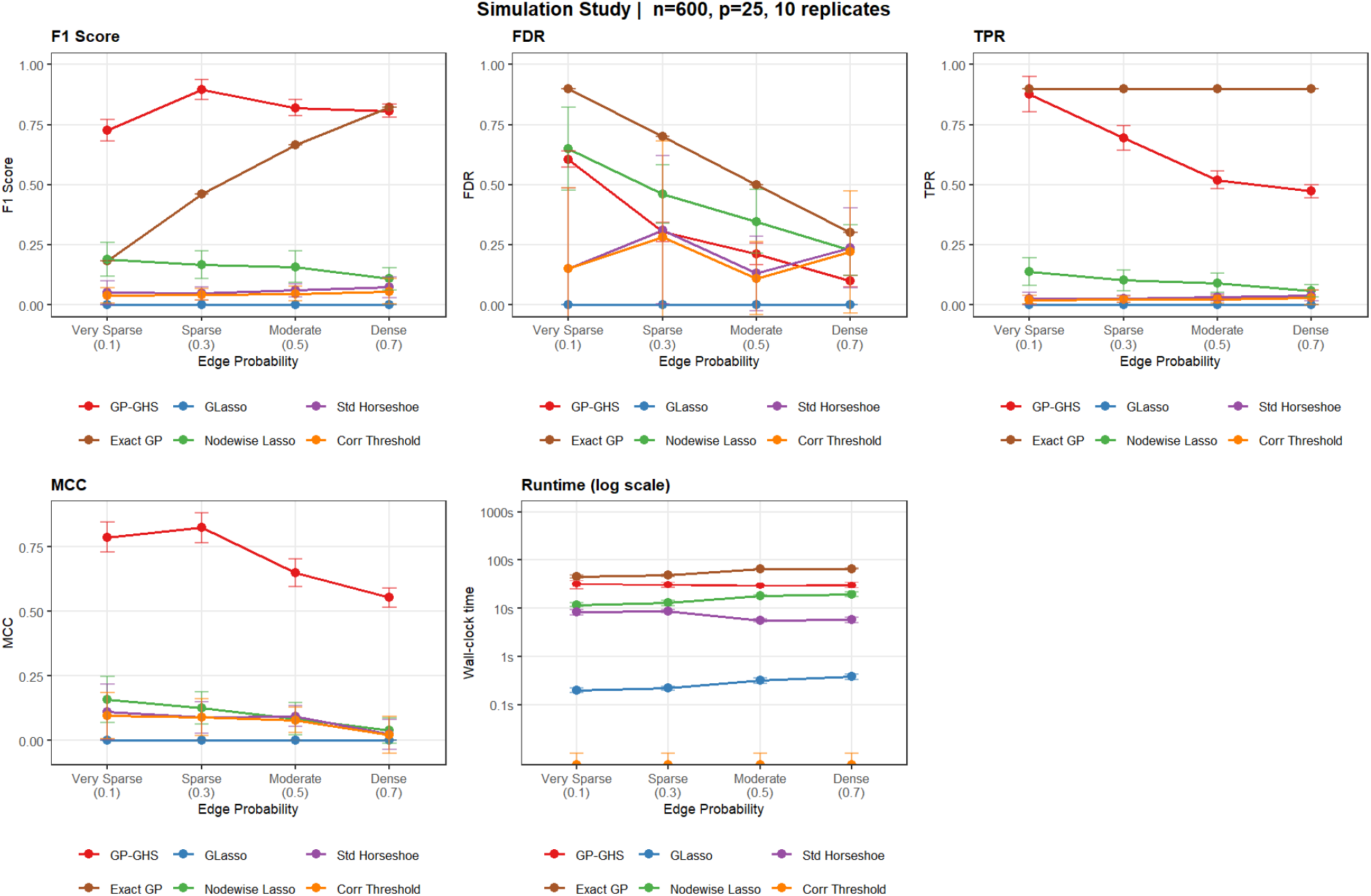
Simulation results for *n* = 600, *p* = 25, scale-free true graph, 10 replicates per sparsity level. Each panel shows mean ± one standard deviation across replicates. Edge probability on the horizontal axis corresponds to expected edge counts of approximately 30 (very sparse, *π* = 0.1), 90 (sparse, *π* = 0.3), 150 (moderate, *π* = 0.5), and 210 (dense, *π* = 0.7) out of 300 possible edges. The runtime panel uses a log_10_ scale.

#### Finding 1: GP-GHS retains dominant F_1_ and MCC performance at *p* = 25, with Exact GP again competitive only in dense graphs

GP-GHS leads all methods on F_1_ and MCC across the full sparsity range, and the magnitude of its advantage over non-spatial competitors is comparable to what was observed at *p* = 15. The F_1_ curve peaks at sparse density and remains strong through moderate and dense settings, with only a modest decline relative to *p* = 15 that is small given the substantially harder candidate edge space. No competitor exceeds a fifth of the GP-GHS F_1_ value at any sparsity level. Exact GP again displays a steeply rising F_1_ trajectory that converges with GP-GHS only at the dense extreme, where the prevalence of true edges is high enough that any high-sensitivity method achieves reasonable precision by default. Across all sparser settings, Exact GP lags GP-GHS substantially on both F_1_ and MCC despite maintaining near-perfect sensitivity throughout, confirming that high TPR alone is insufficient for competitive graph recovery when the null edge set is large.

#### Finding 2: Exact GP achieves uniformly high TPR but its FDR inflation worsens with dimension

Exact GP maintains near-constant and uniformly high TPR across all sparsity levels at *p* = 25, reproducing its *p* = 15 behavior exactly. However, its FDR is notably worse at *p* = 25 than at *p* = 15 across the sparse and moderate settings, with the gap most visible at moderate density where FDR approximately doubles relative to the smaller problem. This worsening is mechanistically transparent: the majority-vote selection rule evaluates each node pair independently without any global calibration, so as the null edge set grows with *p*, expected false inclusions accumulate proportionally. GP-GHS avoids this through the global horseshoe parameter *τ*, which adjusts downward as the null space expands and suppresses additional false inclusions in a data-adaptive way. The value of this joint sparsity regularization therefore scales with *p*, and its advantage over Exact GP becomes more pronounced at higher dimension.

#### Finding 3: GP-GHS FDR control improves with graph density but the very sparse regime remains the hardest setting

The GP-GHS FDR profile follows the same qualitative arc as at *p* = 15: high at very sparse and declining monotonically as the graph densifies, reaching good precision at dense settings. The FDR at the very sparse extreme is modestly elevated compared to *p* = 15, reflecting the larger null edge set at *p* = 25, but the decline across sparsity levels is steeper, so that FDR at moderate and dense settings is comparable across the two problem sizes. TPR follows the expected declining pattern with graph density, for the same reasons described in Section 3.4: as the graph densifies, the global shrinkage parameter stabilizes at a larger value that diffuses discriminative power at the margin. The large replicate-to-replicate variability on all GP-GHS metrics at very sparse confirms that this regime is genuinely difficult regardless of dimension, driven by the small ratio of true signals to candidate edges rather than sampler instability.

#### Finding 4: The ablation argument for group structure strengthens at *p* = 25

Standard Horseshoe continues to fail at edge recovery across all sparsity levels at *p* = 25, selecting a moderate number of edges that bear no coherent relationship to the true graph. The mechanism becomes clearer as *p* grows: each nodewise regression now involves substantially more scalar basis coefficients, and independent scalar shrinkage has no mechanism to couple the coefficients belonging to the same neighbor block into a coherent selection decision. The group prior in GP-GHS makes a single binary selection per neighbor block regardless of the block size, so the number of group-level selection decisions per node is *q* = 24 rather than *qm*^2^ = 384. As *q* increases, the absolute number of spurious scalar-level decisions in SHS grows while the group structure of GP-GHS remains anchored at *q* decisions per node, making the advantage of group shrinkage more visible at higher dimension. GLasso and Nodewise Lasso remain near their *p* = 15 performance floors, confirming that neither penalized likelihood method has any mechanism to exploit spatially heterogeneous interaction structure regardless of problem size.

#### Finding 5: Runtime scaling at *p* = 25 is the dominant practical constraint, with Exact GP now the most expensive method

GP-GHS runtime increases substantially from *p* = 15 to *p* = 25, consistent with the 𝒪 ((*qm*^2^)^2^) scaling of the dominant matrix operations inside the Gibbs sampler. Exact GP is now the most expensive method overall, growing faster than GP-GHS because its cost scales as 𝒪 (*p*(*p* − 1)*n*^3^) — the Cholesky factorization at *n* = 600 is repeated for every ordered node pair, and the number of pairs nearly triples from *p* = 15 to *p* = 25. Nodewise Lasso and SHS also increase noticeably, consistent with their 𝒪 (*nq*) regression costs. GLasso and Correlation Threshold remain negligible. Runtime is flat across sparsity levels for all methods, confirming that the bottleneck is problem dimension rather than graph density. The GP-GHS runtime at *p* = 25 under 11-core parallelization is operationally acceptable for moderate-scale tissue imaging studies; for larger datasets, distributing nodewise regressions across a compute cluster or reducing the HSGP basis dimension from *m* = 4 to *m* = 3 are the natural paths to further speedup.

### 3.6 Cross-Dimensional Comparison

Taken together, the *p* = 15 and *p* = 25 experiments establish several conclusions about the scaling behavior of GP-GHS and its competitors. The qualitative dominance of GP-GHS is preserved across both problem sizes, and the absolute performance losses from increasing *p* are small relative to the growth of the candidate edge space, suggesting that the statistical model is well-specified for spatially structured graphs in this dimension range. The primary cost of increasing *p* is computational rather than statistical, and the practical bottleneck for larger applications is runtime rather than inferential quality.

The behavior of Exact GP across both settings reveals a structural property of independent per-pair selection: high TPR is preserved with dimension but FDR control degrades as the null edge set grows, because there is no global mechanism to recalibrate the selection threshold. GP-GHS resolves this through joint group horseshoe regularization, and the gap between the two spatial methods on FDR widens with *p*. Their convergence on F_1_ at dense graphs is a regime-specific artifact of high edge prevalence rather than evidence of statistical equivalence.

The FDR-TPR tradeoff at the very sparse extreme is essentially stable across both settings for GP-GHS, reflecting a structural property of the horseshoe prior when the true signal fraction is small. Adaptive calibration of the global shrinkage parameter or data-driven thresholding of the edge inclusion criterion are natural directions for improving precision in this regime without sacrificing the high sensitivity that distinguishes GP-GHS from all competitors.

## 4 Data Analysis: Spatial Cell-Cell Interaction Networks in Colorectal Cancer

### 4.1 Data and Preprocessing

We apply the GP-GHS framework to the publicly available CODEX dataset of Schürch et al. (2020), which comprises multiplexed single-cell imaging data from 35 colorectal cancer patients across 140 tissue images spanning two pathologically distinct tumor microenvironments: Crohn’s-like reaction (CLR, *n* = 68 images) and diffuse inflammatory infiltrate (DII, *n* = 72 images). Following upstream phenotyping provided with the dataset, each cell is assigned to one of *K* discrete cell types. We retain cell types present in at least 0.5% of all cells across the full cohort, yielding *K* = 15 types: B cells, CD4^+^ T cells, CD8^+^ T cells, CD163^+^ macrophages, CD68^+^ macrophages, regulatory T cells (Tregs), memory CD4^+^ T cells, generic immune cells, granulocytes, plasma cells, smooth muscle, stroma, tumor cells, vasculature, and adipocytes. Each patient contributes exactly four images, and patients are nested within pathology group, with 17 patients assigned to CLR and 18 to DII.

### 4.2 Spatial Feature Construction via Kernel Density Estimation

The GP-GHS model requires a spatially referenced, continuous-valued feature matrix as input. Raw cell coordinates constitute a marked spatial point process with substantial within-image sparsity, which is unsuitable for direct spatial regression. To convert point-pattern observations into a form amenable to GP-GHS, we estimate a smooth spatial abundance surface for each cell type in each image using two-dimensional kernel density estimation (KDE) to capture the heterogeneity gradient of each cell type through the marginal densities. For each image, we define a regular 30 × 30 grid over the image bounding box, expanded by 5% on each side to mitigate boundary effects, yielding 900 grid locations per image. The KDE for each cell type is evaluated on this common grid using MASS::kde2d, with bandwidth selected independently per spatial axis via the normal reference rule. Cell types with fewer than five observed cells in a given image are assigned a small constant density (10^−6^) to maintain numerical stability. The resulting 900 × *K* density matrix is log-transformed after scaling by 10^4^ to prevent numerical underflow, and each column is subsequently standardized to zero mean and unit variance. The grid coordinates serve directly as spatial locations for the HSGP basis construction. To this end, we use these marginal KDE profiles for individual cell types as cell expressions in our model.

### 4.3 GP-GHS Network Inference

We fit the GP-GHS model independently to each of the 140 images. For each image, *K* = 15 node-wise regressions are executed in parallel, each treating one cell type’s KDE surface as the response and the remaining *K* − 1 surfaces as spatially structured predictors. The HSGP approximation uses *m* = 5 basis functions per spatial dimension (*m*^2^ = 25 total basis functions) with a Matérn-3*/*2 covariance kernel. The MCMC sampler runs for 3,000 iterations with a burn-in of 1,000 and a thinning factor of 5, retaining 400 posterior samples per node. Nodes whose response variance falls below 10^−8^ after standardization are excluded from the regression models and assigned no edges, as a spatially constant KDE surface provides no information for regression. An edge between cell types *s* and *s*^′^ is declared active if the posterior mean of the shrinkage statistic *κ*_*ss*_′ *<* 0.5, where *κ*_*ss*_′ ∈ (0, 1) with values near zero indicating strong evidence of interaction and values near one indicating near-complete shrinkage toward the null. The adjacency matrix is symmetrized via an AND rule, requiring mutual inclusion from both directional regressions, and the *κ* matrix is symmetrized by taking the element-wise minimum, which constitutes the more conservative choice by requiring both nodes to provide evidence of a shared interaction. The entire pipeline ran to completion across all 140 images, with a total wall-clock time of approximately 251 minutes on a multi-core server.

## Results

### Spatial interaction graphs from CRC tissue images

We applied GP-GHS to a colorectal cancer (CRC) tissue imaging dataset comprising 140 images from 35 patients, with four images acquired per patient. Pathology classification assigned 17 patients (68 images) to the CLR group and 18 patients (72 images) to the DII group. Each image contained spatial coordinates and cell type labels across 16 cell types: adipocytes, B cells, CD163+ macrophages, CD4+ T cells, CD68+ macrophages, CD8+ T cells, generic immune cells, granulocytes, memory CD4+ T cells, plasma cells, smooth muscle, stroma, Tregs, tumor cells, and vasculature.

For each image, we first estimated cell type abundance surfaces using two-dimensional kernel density estimation on a 30 × 30 spatial grid, producing a 900 × 16 feature matrix **Y** with one column per cell type. Grid-referenced densities were log-transformed and standardized prior to modeling. GP-GHS was then fit independently to each image, yielding an undirected spatial interaction graph with binary adjacency and a symmetric shrinkage coefficient matrix ***κ***. Under the horseshoe prior convention, *κ*_*ss*_′ ≈ 0 indicates a spatially covarying interaction between cell types *s* and *s*^′^, while *κ*_*ss*_′ ≈ 1 indicates shrinkage toward the null. Symmetrization of directional edge estimates followed an AND-rule for the adjacency matrix and a minimum-rule for *κ*, the latter selecting the directional posterior with stronger evidence of interaction.

### Differential testing of spatial interactions between CLR and DII

To identify edges with differential interaction strength between pathology groups, we tested all 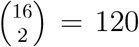 possible cell type pairs; after restricting to edges observed across images, 105 edges were retained for testing. For each edge (*s, s*^′^), we modeled the image-level logit-transformed shrinkage coefficient logit(*κ*_*ss*_′_,*i*_) using a linear mixed model (LMM):

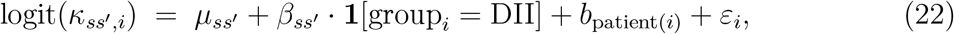

where 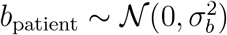 is a patient-level random intercept that accounts for the correlation among the four images per patient, and *ε*_*i*_ ∼ 𝒩 (0, *σ*^2^ ). The reference level was CLR. The fixed effect 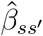 is the primary inferential quantity: it represents the group difference in logit(*κ*) after adjusting for within-patient correlation. The group-specific marginal kappa estimates 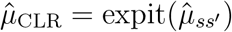 and 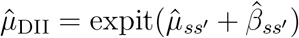 were back-transformed for interpretability. A likelihood ratio test comparing the full model to a null model without the group term produced edge-level *p*-values. Multiple testing correction was applied using the Benjamini–Hochberg procedure at a false discovery rate (FDR) threshold of 5%.

### Differentially active spatial interactions

Thirteen of the 105 tested edges reached statistical significance at FDR *<* 0.05 (Table 1). All 13 significant edges showed stronger spatial co-variation in DII relative to CLR, as indicated by uniformly negative 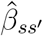 values (Figure 5). No edge exhibited significantly weaker interaction in DII.

**Table 1:** Significant differentially active edges (LMM FDR *<* 0.05), ordered by FDR. 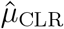 and 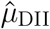 are group-specific marginal kappa estimates back-transformed from the logit scale via the fitted LMM. 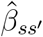 is the LMM fixed effect on the logit scale (DII vs. CLR); all values are negative, indicating stronger spatial interaction in DII. 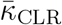 and 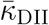 are raw image-level kappa means, retained as diagnostics only.

| Edge | $\hat{\mu}_{\text{CLR}}$ | $\hat{\mu}_{\text{DII}}$ | $\hat{\beta}_{ss'}$ | $\bar{\kappa}_{\text{CLR}}$ | $\bar{\kappa}_{\text{DII}}$ | FDR |
| --- | --- | --- | --- | --- | --- | --- |
| CD68+ macros ↔ Tregs | 0.9966 | 0.6984 | -4.838 | 0.5992 | 0.4298 | 0.0021 |
| memory CD4+ T ↔ Tregs | 0.9931 | 0.5620 | -4.722 | 0.6366 | 0.4417 | 0.0021 |
| CD8+ T cells ↔ Tregs | 0.9938 | 0.6547 | -4.435 | 0.6482 | 0.4821 | 0.0022 |
| granulocytes ↔ Tregs | 0.9944 | 0.7207 | -4.227 | 0.6559 | 0.5454 | 0.0022 |
| stroma ↔ Tregs | 0.9940 | 0.6800 | -4.349 | 0.6750 | 0.5109 | 0.0022 |
| Tregs ↔ tumor cells | 0.9975 | 0.8737 | -4.054 | 0.7617 | 0.7173 | 0.0022 |
| Tregs ↔ vasculature | 0.9951 | 0.6331 | -4.771 | 0.6635 | 0.4630 | 0.0022 |
| CD163+ macros ↔ Tregs | 0.9967 | 0.8610 | -3.885 | 0.7693 | 0.7041 | 0.0033 |
| B cells ↔ Tregs | 0.9990 | 0.9017 | -4.694 | 0.7214 | 0.5006 | 0.0057 |
| smooth muscle ↔ Tregs | 0.9982 | 0.9084 | -4.043 | 0.7314 | 0.6241 | 0.0130 |
| CD68+ macros ↔ memory CD4+ T | 0.9719 | 0.6599 | -2.882 | 0.5923 | 0.4601 | 0.0373 |
| CD68+ macros ↔ stroma | 0.9744 | 0.7452 | -2.566 | 0.6176 | 0.5429 | 0.0373 |
| CD68+ macros ↔ smooth muscle | 0.9929 | 0.8932 | -2.817 | 0.7152 | 0.6274 | 0.0469 |

The strongest and most statistically significant differential signals were concentrated around Tregs. Ten of the 13 significant edges involved Tregs as one partner, spanning interactions with CD68+ macrophages (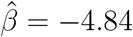, FDR = 0.0021), memory CD4+ T cells (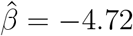, FDR = 0.0021), vasculature (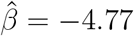, FDR = 0.0022), CD8+ T cells (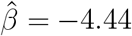, FDR = 0.0022), stroma (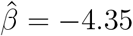, FDR = 0.0022), granulocytes (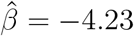, FDR = 0.0022), tumor cells (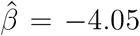, FDR = 0.0022), CD163+ macrophages (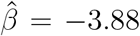, FDR = 0.0033), B cells (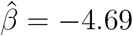, FDR = 0.0057), and smooth muscle (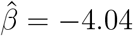, FDR = 0.013). The remaining three significant edges involved CD68+ macrophages interacting with memory CD4+ T cells (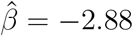, FDR = 0.037), stroma (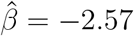, FDR = 0.037), and smooth muscle (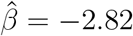, FDR = 0.047).

The LMM-estimated marginal kappa values illustrate the magnitude of these group differences on the kappa scale (Figure 6). In CLR, Treg-involving edges had estimated 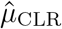 values near 1.0, indicating that these interactions were largely shrunk toward the null, consistent with sparse or spatially diffuse co-localization. In DII, the same edges had 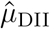 values ranging from approximately 0.56 to 0.90, indicating substantially less shrinkage and thus stronger inferred spatial interaction. For example, the CD68+ macros–Tregs edge had 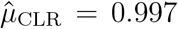 versus 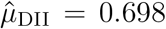, and the memory CD4+ T–Tregs edge had 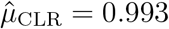 versus 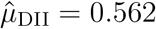.

The beta-hat heatmap (Figure 7) confirms the directional asymmetry across all 105 pairs simultaneously. Edge prevalence heatmaps (Figure 4) corroborate these findings at the graph topology level. The DII group showed visibly higher edge prevalence for Treg-centered interactions and for edges connecting CD68+ macrophages to stromal and immune populations, consistent with the LMM-based differential testing results.

**Figure 4.**
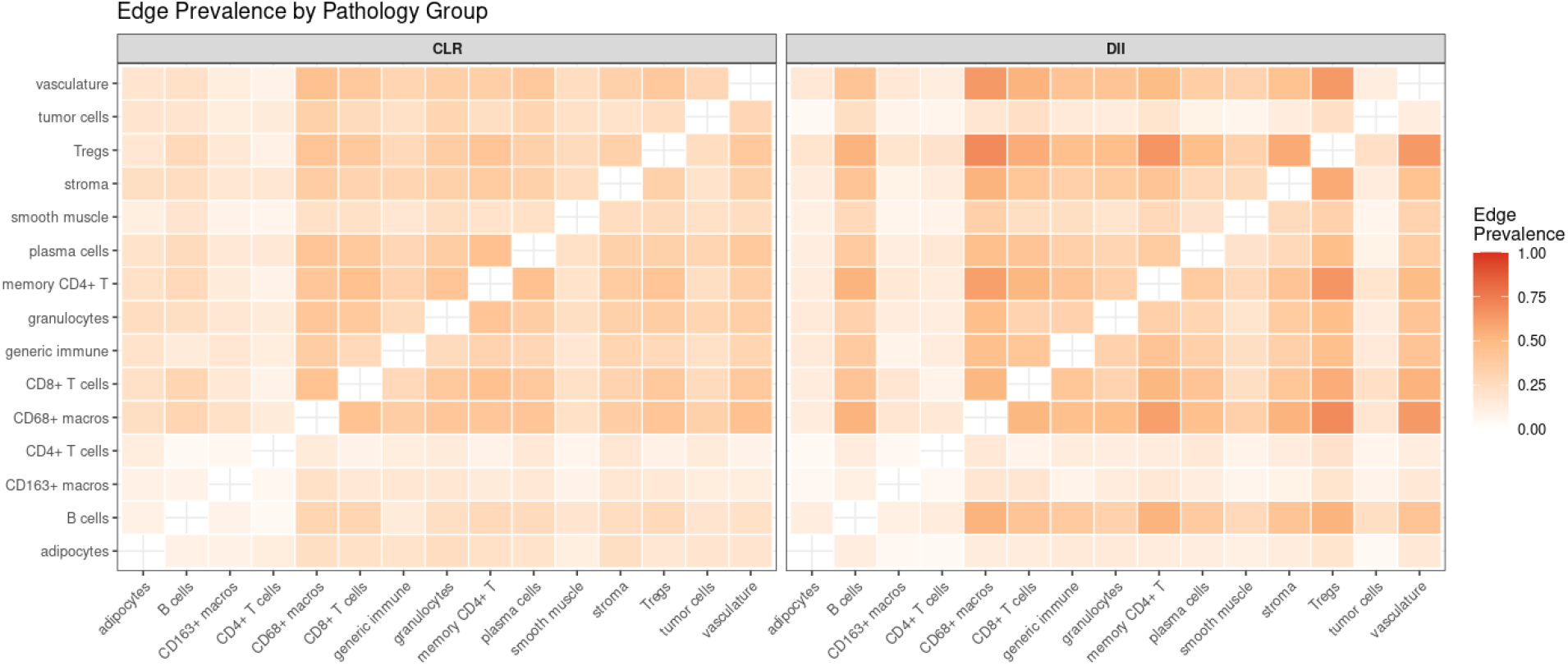
Edge prevalence by pathology group. Each cell (*s, s*^′^) shows the proportion of images within a group in which the GP-GHS AND-rule adjacency assigned an active edge between cell types *s* and *s*^′^. Darker red indicates higher prevalence, meaning the co-localization signal was selected more consistently across images. In CLR, edge prevalence is distributed broadly and at moderate levels across most cell type pairs. In DII, a pronounced cluster of high-prevalence edges emerges around Tregs, with CD68+ macrophages additionally showing elevated connectivity to stromal and immune populations. This topological contrast motivates the formal differential testing carried out in subsequent analyses.

**Figure 5.**
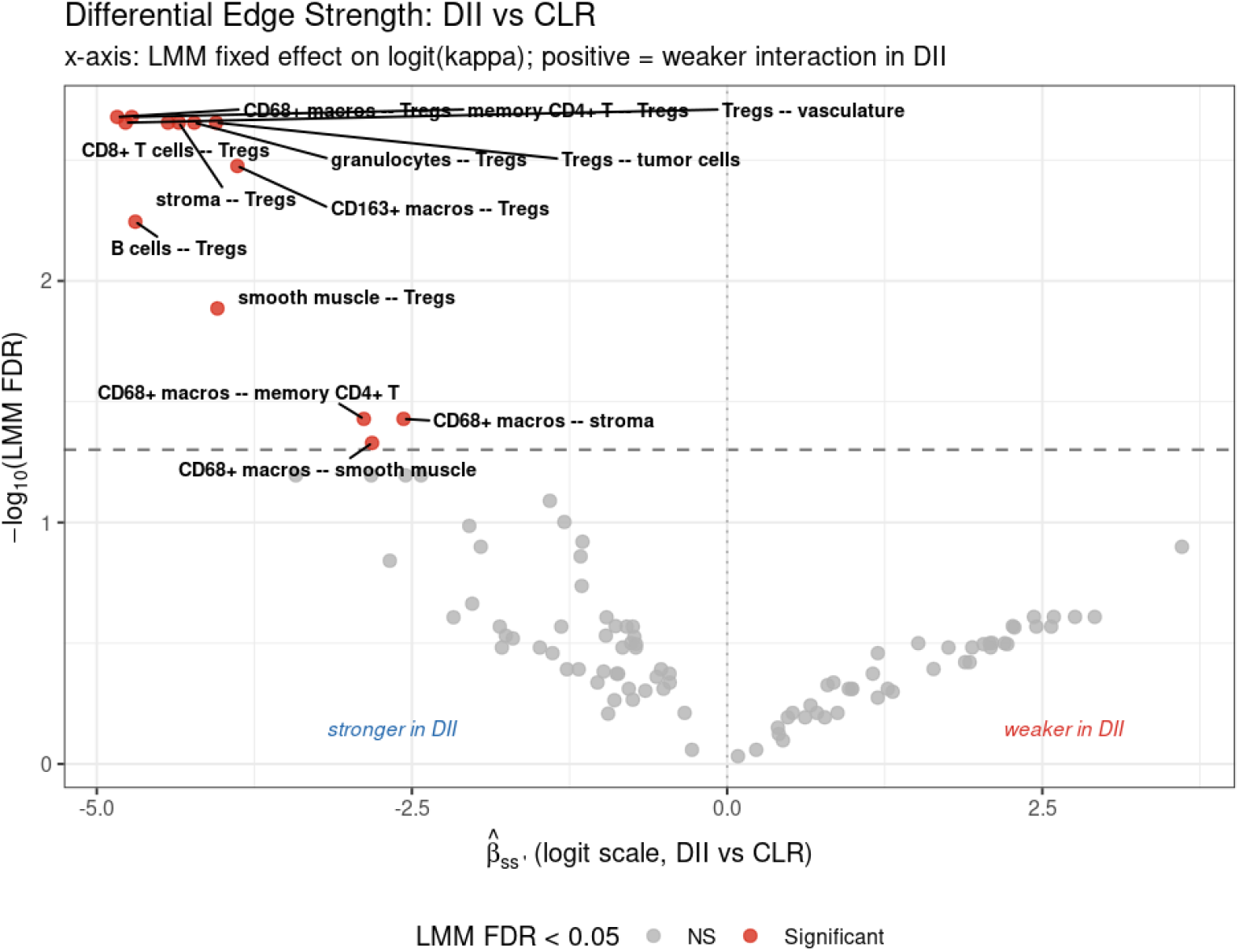
Volcano plot of differential edge strength between DII and CLR. Each point represents one of the 105 tested cell type pair edges. The *x*-axis shows the LMM fixed effect 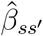 on the logit(*κ*) scale, quantifying the direction and magnitude of the group difference; negative values indicate stronger spatial interaction in DII (lower kappa, less shrinkage) and positive values indicate weaker interaction in DII. The *y*-axis shows − log_10_(FDR), with the dashed horizontal line marking the 5% FDR threshold. Red points are edges significant at FDR *<* 0.05. All 13 significant edges fall on the left side of the axis, confirming that DII is characterized exclusively by stronger, not weaker, spatial co-localization relative to CLR. Labeled edges are dominated by Treg-involving pairs and by CD68+ macrophage interactions with stromal and immune populations.

**Figure 6.**
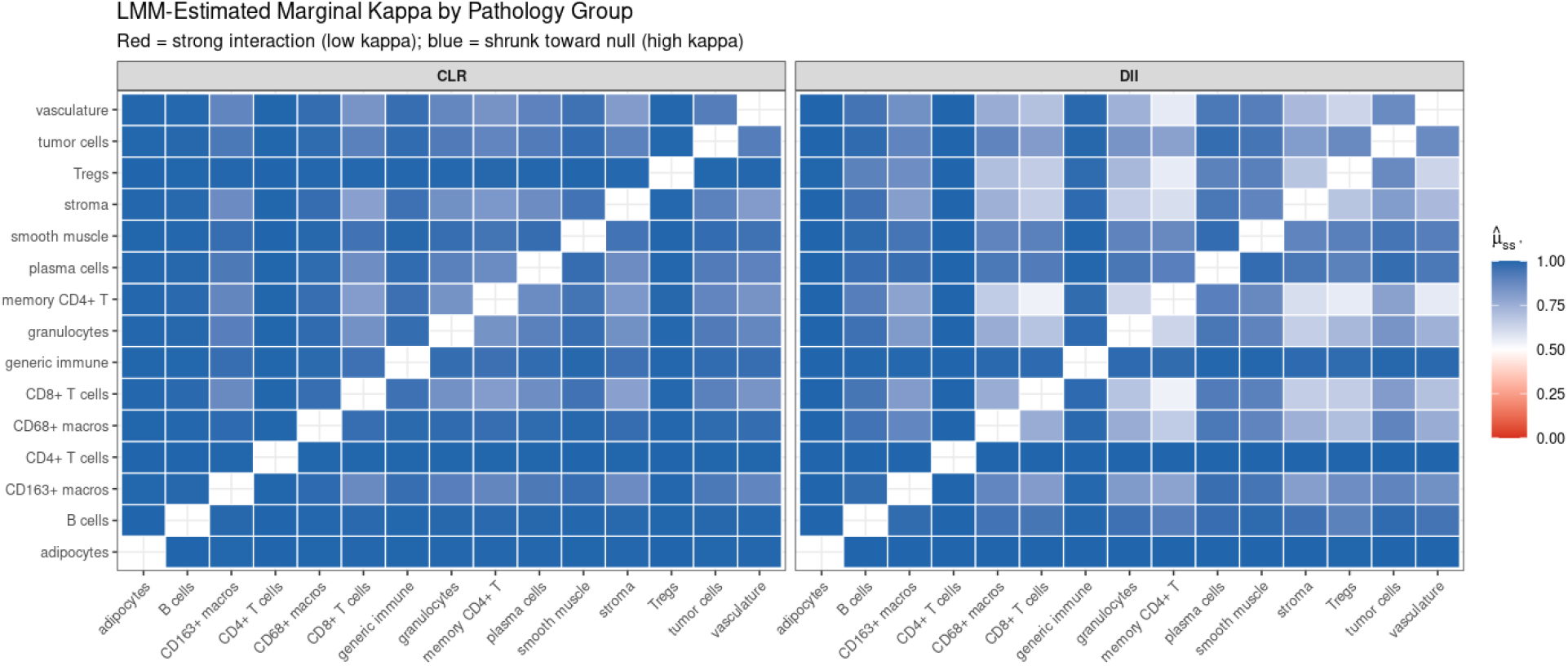
LMM-estimated marginal kappa by pathology group. Each cell (*s, s*^′^) displays 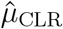 or 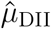, the group-specific marginal kappa back-transformed from the logit scale via the fitted LMM intercept and group effect. Red indicates low kappa (strong spatial interaction) and blue indicates high kappa (shrinkage toward the null). In CLR, nearly all cell type pairs have kappa values close to 1, reflecting a globally sparse interaction structure. In DII, a distinct block of low-kappa entries appears around Tregs and CD68+ macrophages, indicating that these populations are systematically more spatially co-localized with their neighbors in DII tissue. The side-by-side layout enables direct comparison of interaction strength across all pairs simultaneously, with the color contrast between panels visually summarizing the direction and extent of differential rewiring.

**Figure 7.**
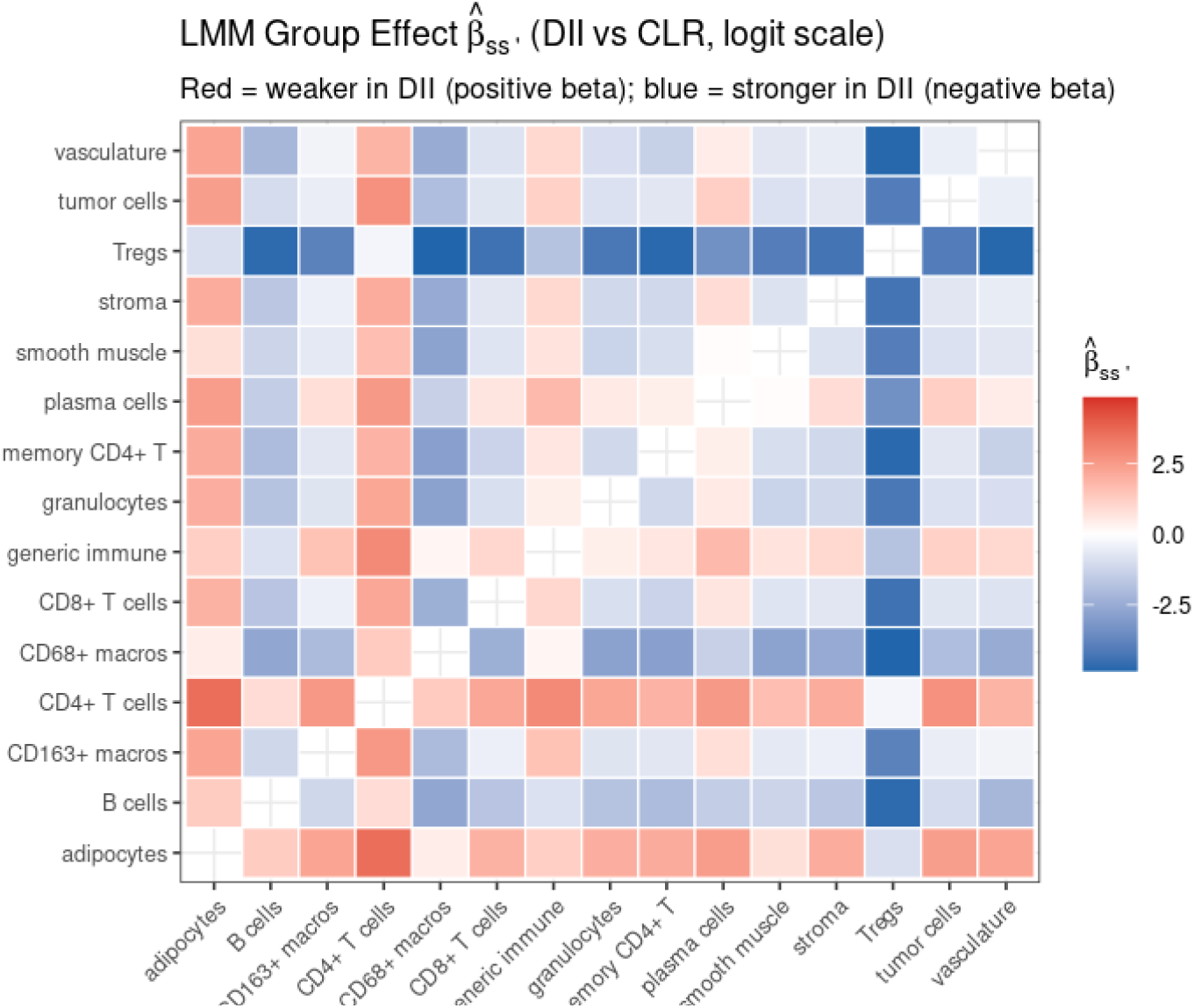
Heatmap of the LMM group effect 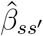 (DII vs. CLR, logit scale). Each cell displays the estimated fixed effect from Equation (22) for that cell type pair, with a symmetric diverging color scale centered at zero. Blue indicates negative 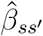, meaning the spatial interaction is stronger in DII; red indicates positive 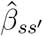, meaning the interaction is weaker in DII. The strongest negative effects cluster in the Tregs row and column and, to a lesser degree, in the CD68+ macrophages row, mirroring the significant edges identified by formal testing. The near-absence of strong positive (red) values across the entire matrix confirms that DII tissue is characterized by a net gain of spatial interactions relative to CLR, rather than a bidirectional rewiring of the cell-cell co-localization network.

#### 4.3.1 Biological Interpretation

The differential testing results point to a Treg-centric immunosuppressive network that is selectively amplified in DII relative to CLR. In DII tissue, Tregs exhibit significantly stronger spatial co-localization with nearly every major immune and stromal compartment represented in the data, including CD68^+^ macrophages, CD163^+^ macrophages, CD8^+^ T cells, memory CD4^+^ T cells, B cells, granulocytes, tumor cells, vasculature, stroma, and smooth muscle. The LMM-estimated kappa values confirm that these differences reflect genuine changes in spatial dependency structure and not simply an increase in edge prevalence: in CLR, these Treg-involving edges are almost entirely shrunk toward the null 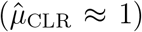, whereas in DII the same edges carry substantially lower posterior shrinkage (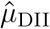 ranging from 0.56 to 0.90), indicating that Treg co-localization with the broader immune microenvironment is a defining spatial feature of DII tumors.

This pattern is consistent with the known biology of the diffuse inflammatory infiltrate subtype. DII tumors are characterized by a heterogeneous, spatially dispersed immune infiltrate in which Tregs are thought to accumulate across multiple tissue compartments and suppress effector immune responses through contact-dependent and cytokine-mediated mechanisms (Schürch et al., 2020). The concurrent spatial coupling of Tregs with both myeloid populations (CD68^+^ and CD163^+^ macrophages) and lymphoid populations (CD8^+^ T cells, memory CD4^+^ T cells, B cells) suggests coordinated immune suppression operating across multiple arms of the adaptive and innate response. The strong Treg ↔ vasculature interaction in DII 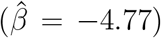 may additionally reflect perivascular Treg niches, a spatial arrangement that has been linked to impaired immune cell trafficking into the tumor parenchyma.

The three significant CD68^+^ macrophage edges↔-with memory CD4^+^ T cells, stroma, and smooth muscle—suggest a secondary axis of spatial reorganization in DII that is independent of Tregs. CD68^+^ macrophages in DII appear to be embedded in a denser stromal and smooth muscle context, which may reflect a myeloid-enriched microenvironment consistent with tumor-associated macrophage polarization toward an immunosuppressive phenotype. The co-localization of CD68^+^ macrophages with memory CD4^+^ T cells in DII could indicate antigen-presentation interactions that are spatially facilitated in this subtype but disrupted or spatially diffuse in CLR.

In contrast, CLR tissue shows a globally sparse spatial interaction structure across all 105 tested edges, with LMM-estimated kappa values near 1 for the majority of cell type pairs. This is consistent with the Crohn’s-like inflammatory architecture of CLR, in which immune infiltration is organized into discrete lymphoid aggregates with a more focal rather than diffuse spatial distribution (Schürch et al., 2020). The relative absence of strong spatial dependencies in CLR may reflect a more compartmentalized immune response, where effector populations are segregated rather than spatially intermingled with regulatory and stromal elements. Importantly, no edges showed significantly stronger interaction in CLR relative to DII, indicating that the network rewiring between subtypes is directionally asymmetric and predominantly reflects interaction gain in DII rather than a bidirectional restructuring of the spatial co-localization network.

Together, these findings demonstrate that GP-GHS recovers interpretable and biologically coherent differences in spatial interaction networks from multiplexed imaging data.

The method identifies not just which edges differ, but quantifies the direction and magnitude of those differences on a well-defined inferential scale, enabling comparisons across pathologically defined patient subgroups at a resolution that standard co-occurrence or neighborhood enrichment analyses do not provide.

## 5 Discussion

We have presented GP-GHS, a Bayesian nodewise regression framework for inferring spatially varying cell-cell interaction networks from multiplexed tissue imaging data. The method combines three components that are each individually motivated and jointly necessary: a Hilbert Space Gaussian Process approximation for scalable spatial field estimation, a group horseshoe prior that enforces edge selection as a group-level binary decision across the full set of spectral basis coefficients, and a nodewise regression strategy that decomposes the *p*-dimensional graphical model problem into *p* parallelizable regressions. The simulation study established that no competitor recovers spatially structured graphs with meaningful accuracy, and the analysis of 140 colorectal cancer tissue images demonstrated that the framework recovers biologically interpretable Treg-centered interaction networks that differ coherently between two pathologically defined patient subgroups.

A natural question is why GP-GHS characterizes only the presence of a cell-cell interaction rather than its sign, magnitude, or spatial pattern. Three properties of the multi-image setting make these summaries difficult to report. First, the sign and magnitude of 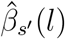 can vary continuously across the tissue within a single image: a co-localization in the tumor core may coincide with spatial exclusion at the invasive margin, so neither a global sign nor a global magnitude is well-defined. Second, there is no common spatial coordinate system across images from different patients: the tissue domain ℒ is image-specific, precluding direct averaging of spatial fields across images. Third, the nodewise regressions are not jointly constrained, so the magnitudes of 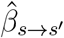 and 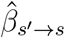 are not on a common scale. Given these constraints, the posterior shrinkage score *κ*-which summarizes how strongly the group prior suppresses the entire interaction field toward zero – is the most portable and interpretable summary across images and patients.

### The group prior is the central modeling contribution

The most important design decision in GP-GHS is the placement of a single local shrinkage parameter *λ*_*j*_ per neighbor block rather than per basis coefficient. The simulation study makes the consequences of this choice concrete. Standard Horseshoe, which shares every other component of the model but applies independent scalar shrinkage to each of the *qm*^2^ basis coefficients, fails to recover any true edges across all sparsity levels and both problem sizes. This failure is not incidental: it reflects the structural mismatch between a prior that makes *m*^2^ independent coefficient-level decisions and a scientific question that requires a single edge-level decision. When *m*^2^ = 16 or 25 basis functions represent a single interaction field, independent shrinkage accumulates *m*^2^ independent opportunities for false activity, which in practice means the posterior for individual coefficients never commits clearly to zero or nonzero. The group prior resolves this by making the *m*^2^ coefficients within a block rise and fall together under a single *λ*_*j*_, so the posterior concentrates sharply at either the all-zero (edge absent) or all-nonzero (edge present) regime. This is precisely the property that produces TPR above 0.85 in the very sparse setting where competitors achieve less than 0.15.

### Limitations and directions for future work

Several limitations of the current framework are worth addressing in future work. First, the KDE-based preprocessing converts a marked point process into a spatial regression input, which involves bandwidth selection and grid resolution choices that could influence the inferred networks. A more principled treatment would model the cell type point process directly, for instance through a log-Gaussian Cox process formulation where the latent intensity surfaces are the regression inputs. This would propagate uncertainty from the point process estimation into the network inference step rather than treating the KDE surfaces as fixed known quantities, which they are not.

Second, as demonstrated with CRC data application, when one has multiple images to compare, the current framework is implemented by fitting one spatial network particular to each image, and then summarizing and comparing the networks across images in a post-hoc linear mixed model. An alternative hierarchical formulation would model all images jointly, placing a prior over the distribution of image-level networks within each pathology group and directly estimating group-level consensus networks with uncertainty quantification. Such a model would borrow strength across images within a group, potentially recovering weaker edges that are present in a majority of images but not strongly enough to survive image-level thresholding. The within-patient correlation structure, which we currently handle only at the testing stage through the random patient effect in the LMM, could be incorporated into the joint model through a patient-level network prior but this increases the computational complexity since we have image-specific GPs.

Third, the computational cost of GP-GHS at *p* = 25 is approximately 31 minutes per image, which becomes a practical bottleneck for whole-slide imaging datasets with hundreds of tissue sections. The dominant cost is the 𝒪 ((*qm*^2^)^2^) matrix operations inside the Gibbs sampler. Two directions for acceleration are worth pursuing: variational inference approximations to the group horseshoe posterior, which would replace the MCMC sampler with a deterministic optimization that scales more favorably, and sparse GP approximations beyond HSGP that exploit the irregular spatial layout of cell locations more efficiently than a regular grid basis.

Fourth, while the HSGP basis functions enforce spatial smoothness through the spectral prior, they do not incorporate any tissue-level structural information such as tissue compartment boundaries or histological annotations. In practice, cell-cell interactions may be qualitatively different in the tumor core, the invasive margin, and the stroma, and a model that can assign separate interaction networks to pre-defined tissue regions, or that learns spatially discontinuous interaction patterns, would be more biologically interpretable Bhadury et al. (2026). Extending the framework to domain-partitioned GPs or spatially nonstationary kernels is a natural direction for future methodological work.

Finally, the current framework does not distinguish between direct cell-cell contact interactions and paracrine signaling interactions that operate over longer spatial scales. The inferred edges reflect spatial conditional dependencies at the scale of the KDE bandwidth and grid resolution, potentially conflating these two mechanistically distinct interaction types. Incorporating spatial scale explicitly, for instance by including multiple bandwidth scales in the KDE or by using a multi-resolution basis expansion, could provide finer-grained decomposition of interaction types.

### Conclusion

GP-GHS addresses a genuine gap in the spatial omics analysis toolkit. Existing graphical model methods either ignore spatial coordinates entirely or treat spatial structure as a nuisance to be residualized rather than a source of signal to be modeled. The group horseshoe prior operating on GP spectral basis coefficients is, to our knowledge, the first method that simultaneously enforces spatial smoothness on each inferred interaction field and makes edge inclusion a coherent group-level decision. The resulting method recovers spatially structured graphs with substantially higher accuracy than all competitors in simulation, and it identifies a biologically coherent Treg-centered immunosuppressive network in colorectal cancer that differentiates between pathologically distinct tumor microenvironments. As multiplexed imaging platforms become standard tools in translational oncology and the number of simultaneously profiled cell types continues to grow, methods that can faithfully represent the spatial heterogeneity of the tumor microenvironment will be essential for translating imaging data into mechanistic hypotheses and ultimately into clinical biomarkers.

## 6 Data and Code Availability

The CODEX dataset is publicly available from Schürch et al. (2020). R package implementation of GP-GHS methodology will be made available at https://github.com/sagnikbhadury/xxxxx with detailed documentation and tutorial examples.

## 7 Author Contributions

S.B., J.G. and A.R. conceived the study and developed the statistical methodology. S.B. implemented the computational framework and contributed to manuscript writing. S.B. contributed to data processing and validation. All authors contributed to manuscript preparation and approved the final version.

## 8 Acknowledgment

During the preparation of this manuscript, the authors used University of Michigan GPT for language editing and sentence-level phrasing suggestions. No AI tool was used to generate results, statistical analyses, proofs, or scientific conclusions. The authors reviewed, verified, and edited all content and take full responsibility for the manuscript.

